# Prediction of lncRNA-protein interacting pairs using LLM embeddings based on evolutionary information

**DOI:** 10.1101/2025.11.10.687101

**Authors:** Shubham Choudhury, Nisha Bajiya, Gajendra P. S. Raghava

**Affiliations:** Department of Computational Biology, Indraprastha Institute of Information Technology, Okhla Phase 3, New Delhi-110020, India

**Author notes:** **Corresponding Author** Prof. Gajendra P. S. Raghava, Professor, Department of Computational Biology, Indraprastha Institute of Information Technology, Delhi, Okhla Industrial Estate, Phase III, New Delhi, India – 110020, Web: http://webs.iiitd.edu.in/raghava/. **Mailing Address of Authors** Shubham Choudhury (SC), Nisha Bajiya (NB).

**Keywords:** lncRNA–protein interaction, Convolutional neural network (CNN), DNABERT-2, ESM-2-t32, Evolutionary information, Embeddings

## Abstract

Interactions of long non-coding RNAs (lncRNAs) with proteins is responsible for numerous cellular processes, including transcriptional regulation, chromatin remodeling, cell differentiation, and intracellular signaling. In the past, numerous computational methods have been developed for predicting lncRNA–protein interacting (LPI) pairs. This study describes a highly accurate and reliable method for predicting LPI pairs built on largest possible non-redundant dataset having 262,244 interacting and equal number of non-interacting pairs. Initially, similarity-based approach BLAST has been tried which have poor discriminative power, due to low sequence similarity. Subsequently, we developed CNN based models and machine learning based models using traditional features and embedding. Our CatBoost model developed using embedding generated by DNABERT-2 and ESM-2-t30 achieved AUC of 0.989 with MCC 0.915 on an independent dataset. Our method performs better than existing methods on an independent dataset. We developed standalone software and web server lncrnaPI for predicting LPI pairs, scanning lncRNA interacting proteins in proteome and protein interacting lncRNA in genomes (https://webs.iiitd.edu.in/raghava/lncrnapi/).

**HIGHLIGHTS:**

- Discrimination of LncRNA-protein interacting and non-interacting pairs.
- Non-redundant dataset of 262,244 interacting and 262,244 non-interacting pairs.
- Embedding of LncRNA using DNABERT-2 and protein using ESM-2-t30.
- Rapid scanning of lncRNA interacting proteins at genome scale.
- A web server and software for predicting LncRNA-protein interacting pairs.

## Introduction

Long non-coding RNAs (lncRNAs) are defined as a class of RNA transcripts exceeding 200 nucleotides with limited or no protein-coding potential. Their regulatory functional roles are mainly mediated through their interactions with proteins, resulting in the formation of ribonucleoprotein (RNP) complexes. For instance, the lncRNA XIST controls X-chromosome inactivation by the recruitment of the Polycomb Repressive Complex 2 (PRC2)^1^, whereas MALAT1 modulates alternative splicing through its association with serine/arginine-rich (SR) proteins^2^. Such LPIs are crucial to cellular homeostasis, and they also modulate transcriptional regulation, chromatin remodeling, and the structural organization of subcellular compartments. The dysregulation of these interactions has been implicated in the pathogenesis of numerous human diseases, such as cancer and neurodegenerative disorders^3^. Understanding of the lncRNA-protein interactome is critical for improving fundamental cell biology research and developing new therapeutic techniques. The identification of LPIs has relied heavily upon high-throughput experimental techniques - such as RNA immunoprecipitation (RIP) and cross-linking immunoprecipitation (CLIP) variants, including PAR-CLIP^4^ and iCLIP^5^. Although these methods provide an accurate picture of the RNA-protein landscape, they still suffer from significant drawbacks. These include their labor-intensive nature, high costs, and need for cellular material.

To overcome above limitations, the scientific community has developed computational methods to scan interacting pairs at genome scale. Early computational approaches were predicated by training classical machine learning classifiers using features derived from RNA and protein sequences. Methodologies such as lncPro^6^ and RPI-Pred^7^ employed features like k-mer frequency, secondary structure, and physicochemical properties as input for classifiers such as Support Vector Machines (SVMs) and Random Forests. In recognition of the possibility that these handcrafted features might not fully encapsulate the biological context, subsequent methodological advancements incorporated network information. For instance, LPIHN^8^ utilized a random walk with restart algorithm on heterogeneous networks constructed from lncRNA expression similarity and protein-protein interaction data. The next generation of LPI prediction tools - LPI-BLS, RPI-SE, LPI-HyADBS and SAWRPI, were based on ensemble strategies where outputs from multiple models have been combined to improved prediction accuracy^9–12^.

Recently, a paradigm shift has occurred toward advanced deep learning architectures that includes Graph Convolutional Networks (GCNs), Graph Autoencoder^13,14^, and Graph Transformers^15^. Comprehensive details about the 32 existing tools have been provided in Supplementary Table S1.

One major limitation of existing methods is that most models have been trained on small and outdated datasets, primarily derived from older versions of the NPInter v2.0 database^16^. Except for a few methods such as LPI-BLS^17^, many existing tools are not publicly available for predicting LPI pairs. To address these limitations, we systematically developed a highly accurate and reliable approach for LPI prediction. Our models were trained, tested, and evaluated on the largest available dataset curated from the latest release of NPInter. We explored both alignment-based and alignment-free strategies. For alignment-based prediction, we implemented the widely used BLAST tool. For alignment-free approaches, we built machine learning and deep learning models using sequential features and evolutionary information. Recent advances in large language model (LLM) embeddings have showed strong potential and therefore, we also developed LLM-based machine learning models. To support the academic community, we provide both a web server and standalone software for predicting LPI pairs and scanning genomes for potential lncRNA–protein interactions.

## Materials and Methods

### Dataset Curation

In this study, we obtained LPI pairs from NPInter v5.0, which contains 2,596,183 experimentally validated non-coding RNA (ncRNA) and biomolecule interactions^18^. To create a clean non-redundant dataset, we implemented a multi-stage filtering protocol. Initially, the database was filtered to retain relevant interacting pairs. We only consider lncRNA-Protein interacting pairs in Homo sapiens. This initial step yielded a total of 488,982 LPI pairs. We filtered lncRNA sequences to include only those with a length between 200 and 20,000 nucleotides. For the protein sequences, we removed any entries containing non-standard amino acids (U, O, B, Z, J, X), which could interfere with downstream feature extraction. The dataset was reduced to 370,242 interaction pairs after this filtering process.

We deployed the CD-HIT tool to cluster lncRNA sequences at an 80% identity threshold and protein sequences at a 95% identity threshold^19^. Post redundancy removal, a final positive dataset of 262,244 unique LPIs was generated. We generated negative samples by scrambling the interacting pairs from our curated positive set. This was achieved by randomly shuffling the protein partners for each lncRNA, ensuring that the resulting pairs were not present in the original positive dataset. An equal number of negative interaction pairs were generated, creating a balanced dataset having a total of 524488 pairs. Finally, the dataset was split into two parts - a training dataset containing 80% of the original dataset (422368 pairs) and a validation dataset containing 20% of the original dataset (102120 pairs). The entire dataset creation process is graphically depicted in Figure 1.

**Figure 1.**
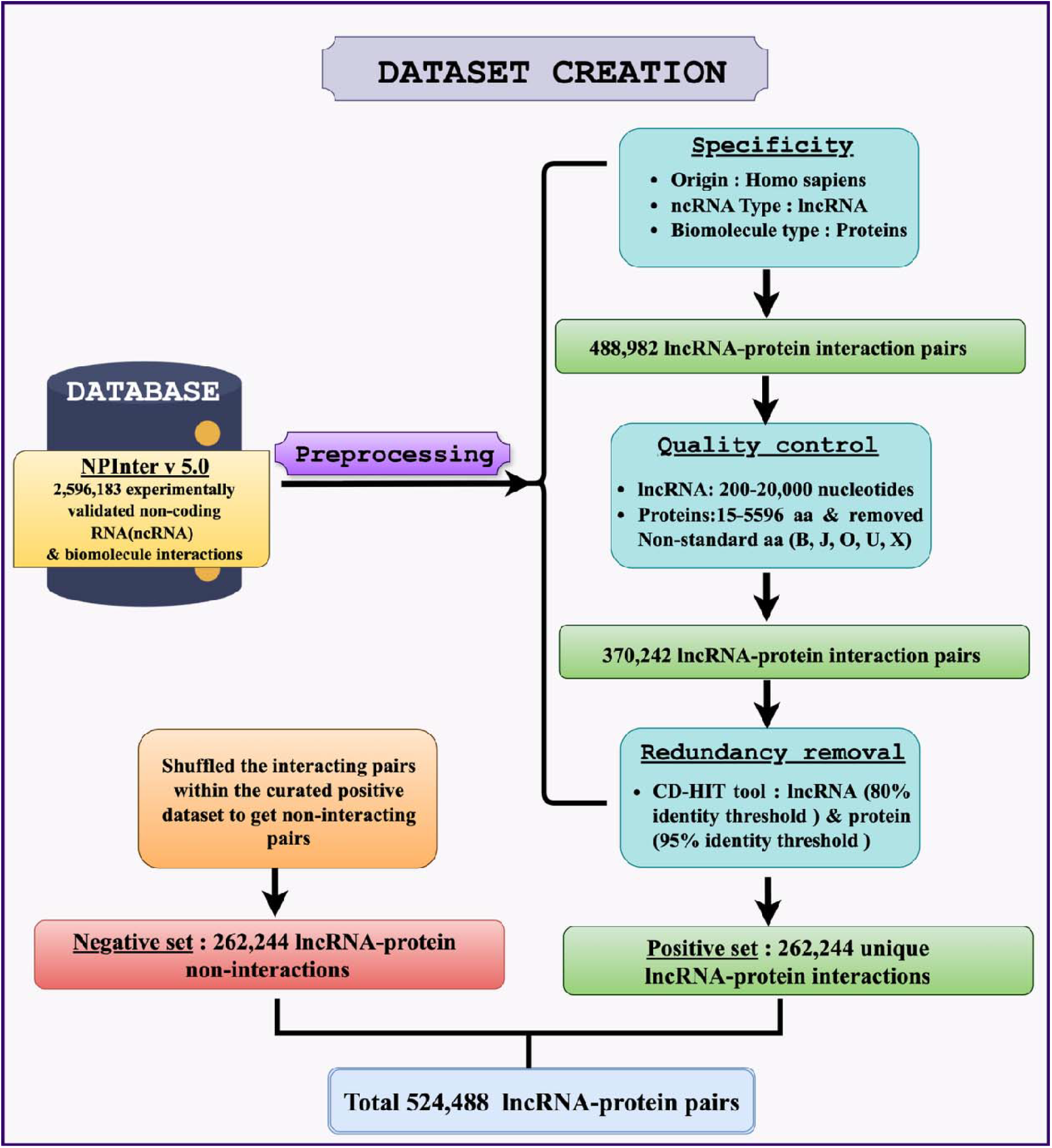
Figure summarizing the dataset creation process used in lncrnaPI.

### Feature Generation

We generated a comprehensive and diverse feature set that includes traditional statistical features, compositional features and evolutionary information, and state-of-the-art contextual embeddings derived from large language models (LLMs). In this study, we compute traditional sequence-based features using the Nfeature and Pfeature toolkits. We generated composition-based descriptors such as k-mer frequencies for lncRNAs and amino acid/dipeptide composition for proteins^20,21^. Correlation-based features were also extracted for lncRNAs via Nfeature, which capture the distribution patterns of physicochemical properties within the nucleotide sequence. The detailed information about the composition and correlation-based features are provided in Supplementary Tables S2 and S3, respectively. For proteins, we have also used evolutionary conservation information as features, by generating Position-Specific Scoring Matrices (PSSMs) using the POSSUM toolkit, which internally uses PSI-BLAST by querying sequences against the SwissProt database^22–24^. The PSSM for each protein provides a probabilistic representation of amino acid substitutions at each position, offering deep evolutionary insights.

In order to deploy Convolutional Neural Networks (CNNs) based models, one-hot encoding was performed on the sequence data. Each lncRNA and protein sequence was converted into a numerical matrix by representing each nucleotide or amino acid as a sparse binary vector. These matrices were then padded or truncated to a fixed length of 2500 for lncRNAs and 1500 for proteins, ensuring uniform input dimensions for the models. Finally, using large language models for lncRNA and proteins, we generated informative contextual embeddings. For lncRNA sequences, we utilized DNABERT-2, a bidirectional transformer model pre-trained on extensive DNA sequence data^25^. For protein sequences, we extracted embeddings from three variants of the Evolutionary Scale Modeling (ESM2) model: ESM2-t30-650M-UR50D, ESM2-t33-3B-UR50D, and ESM2-t48-15B-UR50D ^26^.

### Alignment-based Approach

We developed an alignment-based approach to serve as a baseline method for LPI prediction, as this method utilizes sequence similarity to identify interacting pairs. First, we constructed two separate BLAST databases, one for the lncRNA sequences and one for the protein sequences, using our training dataset. The lncRNA and protein sequences from the validation dataset were then used as queries against their respective BLAST databases. We used two separate strategies for predicting the interacting pairs. The first strategy was a ‘best match’ approach, where we identified the lncRNA and protein pair in the training database with the highest sequence similarity to the queried pair. The interaction label (positive or negative) of this single best-matching pair was then directly assigned to the query pair. The second strategy was a ‘voting model’. In this approach, we considered all significant hits returned by the BLAST search. The final prediction for the query pair was determined by a majority vote, where the label that appeared most frequently among the identified hits was assigned. These alignment-based methods allowed us to assess the extent to which sequence homology alone can predict LPIs within our dataset.

### Machine Learning Approaches

In this study, we used a wide array of machine learning techniques for developing prediction models that include Random Forest, Extra Trees, XGBoost, and CatBoost^27–30^. In case of the Random Forest and Extra Trees models, we configured them with 100 estimators (n_estimators=100). For reproducibility of our results, a random_state of 42 was set for all applicable models. We specifically selected models that support parallel processing, as single-core machine learning models were very time consuming for our large datasets. The XGBoost classifier was implemented with eval_metric=‘logloss’ to guide the gradient boosting process, and the CatBoost classifier was run with verbose=0 to minimize logging output.

### Deep Learning-based Approach

A deep learning framework, designed as a Siamese-inspired Convolutional Neural Network (CNN) architecture, was used to generate one-hot encoded vectors and predict interactions. The model consists of two parallel convolution branches – one for lncRNA and the other for proteins. The lncRNA branch accepts input tensors of shape (2500, 4) and processes them through two successive convolutional blocks. The first block applies a one-dimensional convolutional layer with 128 filters (kernel size = 16), followed by max-pooling (pool size = 4) and dropout (rate = 0.4). The second block uses 256 filters (kernel size = 10) with the same pooling and dropout configurations. The protein branch follows a similar architecture, accepting inputs of shape (1500, 20). Its first convolutional layer contains 128 filters (kernel size = 10), while the second employs 256 filters (kernel size = 8), both followed by identical pooling and dropout layers.

The outputs from the final dropout layers of both branches are flattened and concatenated to form a unified feature vector. This merged representation is then passed through a classification head comprising two fully connected layers with 256 and 128 neurons, respectively, both employing ReLU activation. To reduce the risk of overfitting, a dropout layer (rate = 0.5) is applied after the first dense layer. The final prediction is generated through a single output neuron with a sigmoid activation function, enabling binary classification. The model was optimized using the Adam optimizer (learning rate = 0.001) with binary_crossentropy as the loss function and the Area Under the Curve (AUC) as the primary performance metric. A schematic overview of the model architecture is provided in Figure 2.

**Figure 2.**
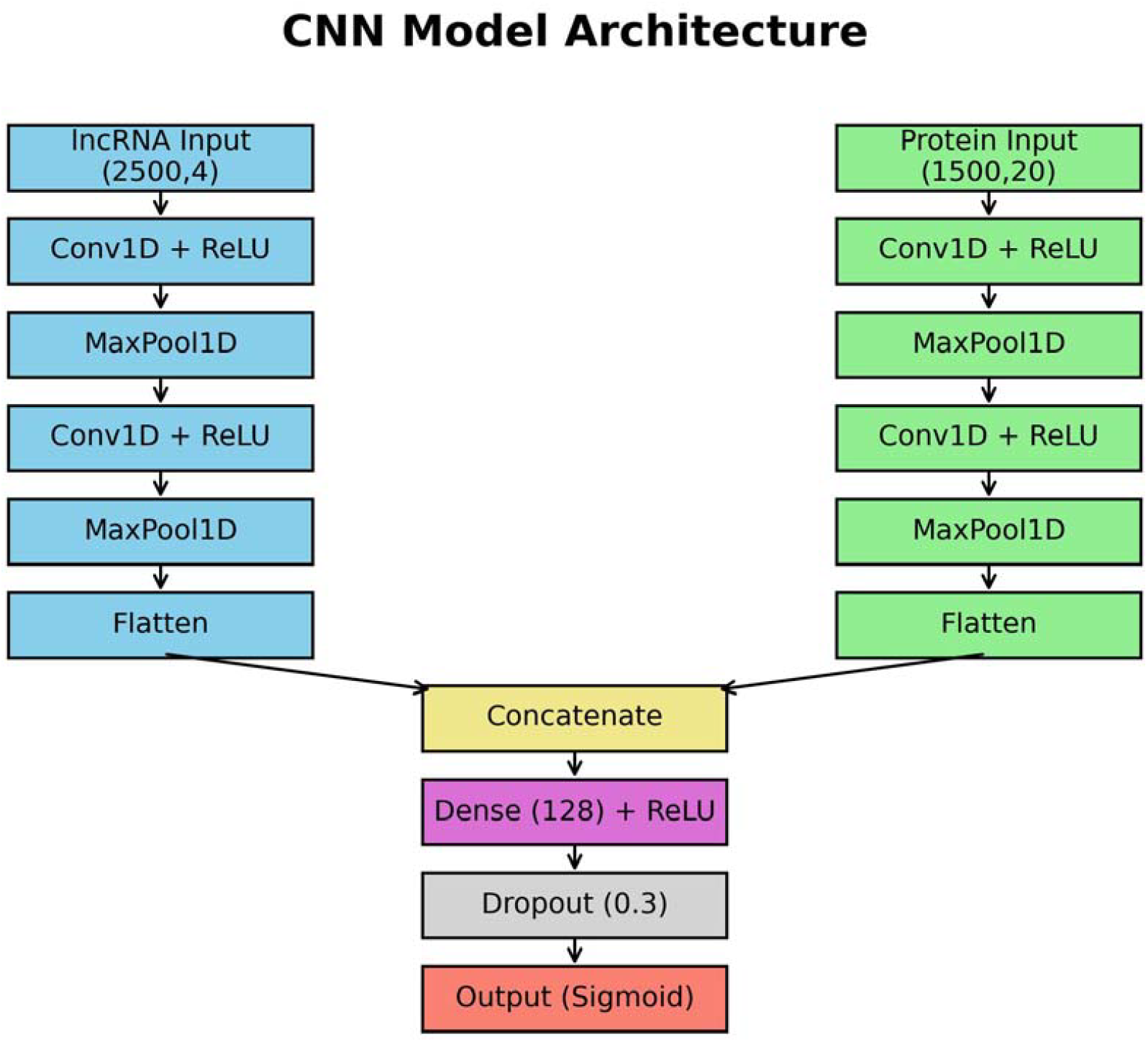
Model Architecture for the CNN model used in lncrnaPI.

### Model Evaluation and Performance Metrics

The predictive performance of our models was evaluated using a comprehensive set of standard statistical metrics derived from counts of true positives (TP), true negatives (TN), false positives (FP), and false negatives (FN). We calculated Sensitivity and Specificity to measure the ability to correctly identify positive and negative interactions, respectively, along with overall Accuracy. The Matthews Correlation Coefficient (MCC) was included as a robust metric that provides a balanced measure of performance. Additionally, we computed the Area Under the Receiver Operating Characteristic Curve (AUC) for a threshold-independent assessment of the model’s overall discriminative power. Models were trained using a 5-fold cross-validation strategy, and the training metrics were calculated as an average over the 5 folds. The validation performance was reported on a held-out independent validation dataset.

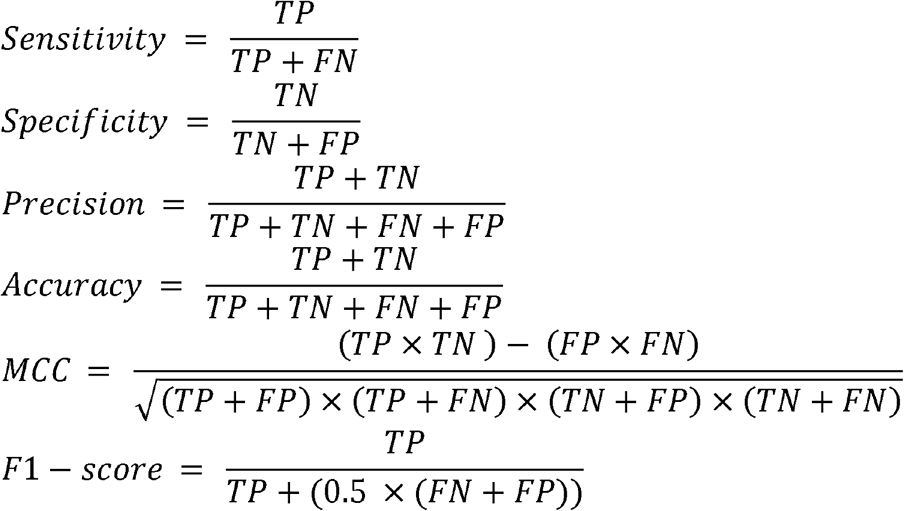

## Results

We systematically evaluated the traditional machine learning classifiers, the deep learning model, and the alignment-based methods using standard evaluation metrics to determine their predictive ability. Figure 3 represents the overall methodology used in this method.

**Figure 3.**
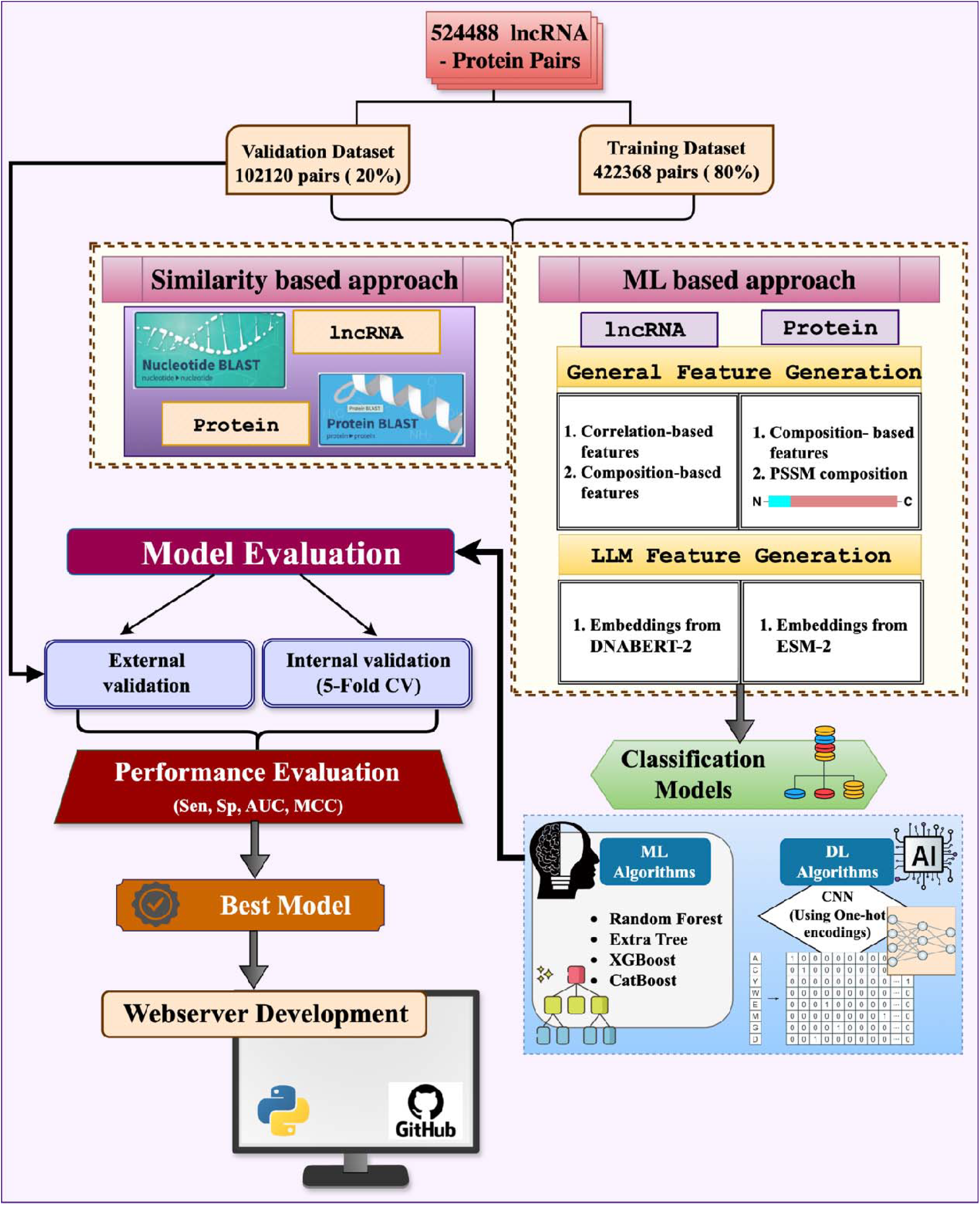
A graphical representation of the overall methodology used in lncrnaPI.

### Compositional Analysis

We performed a detailed compositional analysis of both amino acids and nucleotides to identify which amino acids/nucleotides drive LPIs. We observed that certain amino acids were significantly enriched/depleted in the interacting pairs compared to the non-interacting pairs, as shown in Table 1. Leucine was highly depleted in the interacting pairs and upon using Leucine composition to classify interacting pairs, we achieved accuracy of 71.6%. Conversely, Glycine was enriched in the interacting pairs and Glycine composition could achieve a predictive accuracy of 64.9%. On the other hand, the analysis of nucleotide composition in the lncRNA sequences revealed marginal differences between positive and negative sets, as shown in Table 2, with predictive accuracies lying in the range of 50-51%. This suggests that while simple nucleotide frequency is not a strong driver of interaction specificity, the amino acid composition of the protein partner drives the potential to determine interaction.

**Table 1.** Amino acid compositional of protein involved in interaction pairs and non-interacting pairs and accuracy of threshold-based methods using composition of amino acids.

|  | Percent Composition |  |  |
| --- | --- | --- | --- |
| Amino Acid | Interacting Proteins | Non-interacting Proteins | Predictive Accuracy |
| Depletion (Interacting pairs ≤ non-interacting pairs) | | | |
| L | 6.34 | 9.33 | 0.716 |
| C | 1.2 | 1.9 | 0.612 |
| W | 0.75 | 1.14 | 0.6 |
| T | 4.34 | 4.93 | 0.574 |
| V | 5.05 | 5.93 | 0.572 |
| D | 5.02 | 5.07 | 0.558 |
| A | 6.36 | 7.41 | 0.557 |
| Enriched (Interacting proteins ≥ non-interacting proteins) | | | |
| G | 11.55 | 7.3 | 0.649 |
| Q | 5.78 | 4.69 | 0.61 |
| Y | 3.37 | 2.67 | 0.596 |
| N | 3.92 | 3.33 | 0.592 |
| R | 6.93 | 6.48 | 0.545 |

**Table 2.** Nucleotide Compositional Analysis of lncRNA Sequences.

| Nucleotide | % Composition in Interaction | % Composition in Non-Interaction | Enrichment in Interaction | Predictive Accuracy |
| --- | --- | --- | --- | --- |
| T | 26.05 | 26.04 | Enriched | 0.51 |
| A | 27.29 | 27.57 | Depleted | 0.51 |
| C | 23.29 | 23.1 | Enriched | 0.506 |
| G | 23.37 | 23.29 | Enriched | 0.5 |

### Performance of Alignment-Based Method

Initially, we investigated alignment-based approaches to determine if direct sequence homology could be effectively leveraged for prediction. A key finding was that a large majority of the 102,120 pairs in the validation set did not return any significant BLAST hits from the training database, indicating a high degree of sequence dissimilarity between the training and validation sets. For the subset of pairs where hits were found, the ‘Best Match’ approach found a match for 17,777 pairs and correctly predicted the interaction status for 13,630 of them, achieving an accuracy of 76.67%. The ‘Voting-based’ model, which considers multiple hits, performed slightly better. It found hits for 17,136 pairs and correctly classified 13,659 of them, resulting in a higher accuracy of 79.71%. While these results show that sequence homology has some predictive value, the inability to generate predictions for most pairs and the significantly lower accuracy compared to our ML and DL models demonstrated the limitations of using alignment-based methods as a primary predictive tool in this context.

### Machine Learning based Models

We developed models using machine learning techniques CatBoost, Random Forest, Extra Trees, and XGBoost. These models were developed using different types of features of lncRNA and proteins. In most of the cases CatBoost perform better than other machine learning techniques. The performance of CatBoost method developed using different type features is shown in Table 3, establishing itself as the superior machine learning framework for this task. Our CatBoost model developed using traditional features achieved maximum AUC 0.988. In addition to traditional models, we used DNABERT-2 for computing features or embedding of lncRNA and ESM2 models for computing embeddings of proteins. Our CatBoost model developed using embeddings achieved maximum AUC of 0.989. The summary of the top performing models is provided in Table 3 and the detailed performance results are provided in Supplementary Tables S4 and S5.

**Table 3.** The performance of CatBoost models developed using different types of lncRNA and Protein features.

| lncRNA Features | Protein Features | Training |  |  |  | Validation |  |  |  |
| --- | --- | --- | --- | --- | --- | --- | --- | --- | --- |
|  |  | Accuracy | F1 Score | MCC | AUC | Accuracy | F1 Score | MCC | AUC |
| ALL_COMP | PSSM | 0.953 | 0.954 | 0.907 | 0.987 | 0.953 | 0.953 | 0.907 | 0.987 |
| DACC | APAAC | 0.954 | 0.956 | 0.91 | 0.988 | 0.954 | 0.954 | 0.91 | 0.988 |
| DACC | PSSM | 0.953 | 0.955 | 0.907 | 0.987 | 0.953 | 0.954 | 0.908 | 0.988 |
| DACC | SPC | 0.954 | 0.956 | 0.91 | 0.988 | 0.955 | 0.955 | 0.911 | 0.988 |
| DNABERT-2 | ESM-2 t30 | 0.929 | 0.958 | 0.914 | 0.989 | 0.932 | 0.957 | 0.915 | 0.989 |

<sup>#</sup> DACC - Dinucleotide based auto cross correlation; SPC - Shannon Entropy of Physicochemical Property; APAAC - Amphiphilic Pseudo Amino-acid Composition; PSSM - Position-Specific Scoring Matrix; ESM-2 t30 - esm2\_t30\_150M\_UR50D

### Deep Learning based Model

The performance of our deep learning model, a Siamese-like two-tower CNN trained directly on one-hot encoded vectors, was highly robust and demonstrated the power of end-to-end learning for this prediction task. On the independent validation set, the model achieved a highly competitive Matthews Correlation Coefficient (MCC) of 0.908 and an Area Under the Curve (AUC) of 0.987. These results are particularly noteworthy as they place CNN’s predictive power nearly on par with our best-performing, feature-based CatBoost models. Even though CatBoost models used with simple features or LLM embedding show better performance, the CNN model’s performance has significance as it eliminates the need for feature engineering. The model can generate its own features and learn discriminatory motifs from the dataset directly. The detailed performance metrics for the CNN model is provided in Supplementary Table S6.

### Comparison with Existing Tools

In order to compare our method with the existing tools, we identified 32 existing tools that were designed for LPI prediction. It was seen that most of the tools were either non-functional, inaccessible, or did not provide a standalone version where users could predict using their own data. Only the tool LPI-BLS, developed by Fan et al., allows users to make predictions on their own data. Our method was benchmarked against LPI-BLS using the validation dataset. LPI-BLS exhibited significantly low performances, managing to achieve an accuracy of 0.513, an MCC of −0.003, and an AUC of 0.512. LPI-BLS classified almost all the pairs as non-interacting, as it reported a low sensitivity of 0.012. Figure 4 highlights the performance of our best models with LPI-BLS. The detailed performance metrics are provided in Supplementary Table S7.

**Figure 4.**
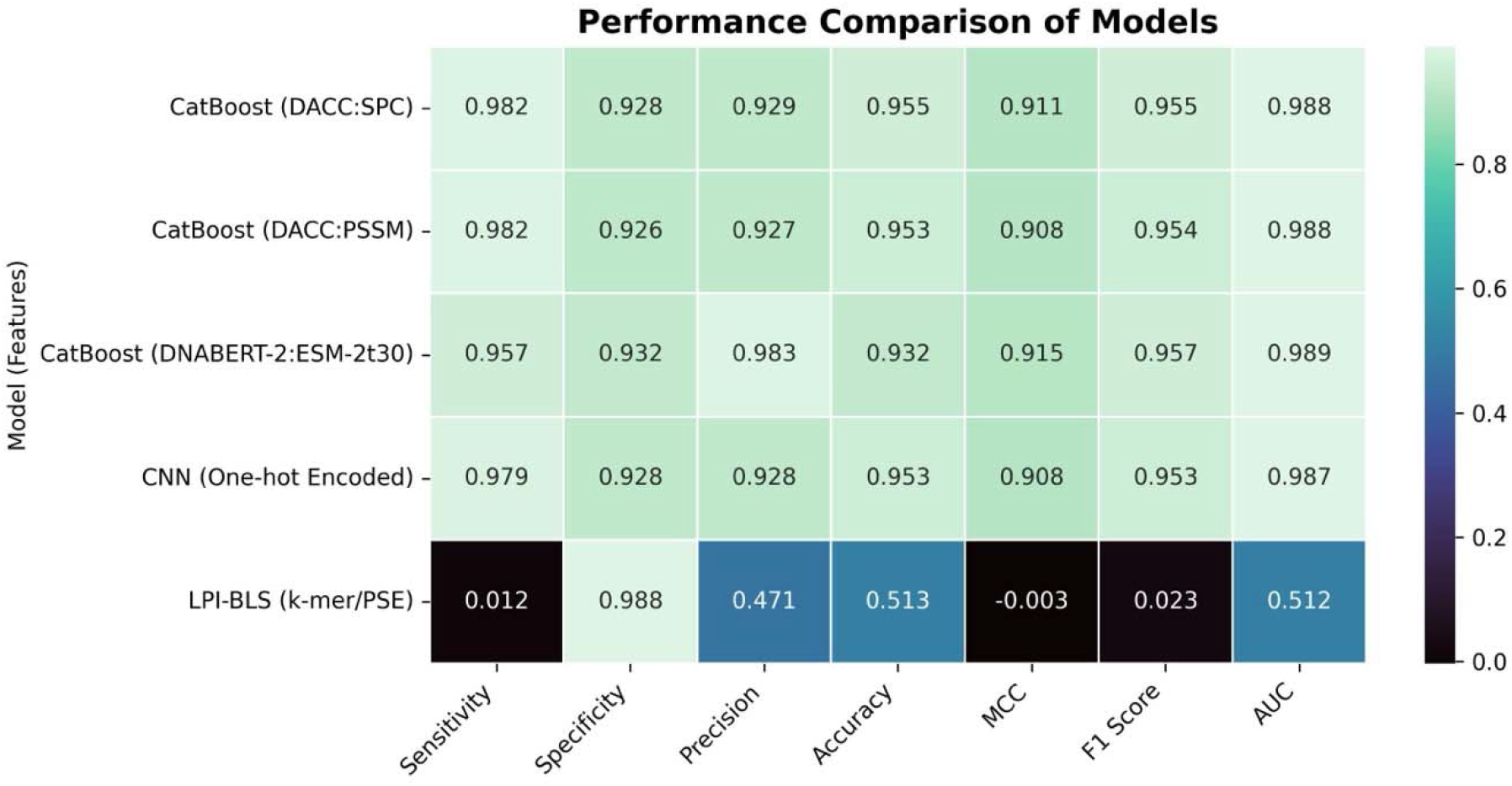
Comparing the performance of LPI-BLS with the other algorithms used in lncrnaPI.

### Rapid Scanning of Interacting pairs

The best performing models in this study rely on features generate using LLMs or features generated from Pfeature/Nfeature. However, these processes can be very time consuming and computationally intensive, when the prediction task involves large proteomes or transcriptomes. In order to fasten up the process, we developed a simple yet fast method for generating features without compromising on predictive accuracy. We observed that a feature-set comprising of composition of 10 amino acids (G, Q, Y, N, R, L, C, W, R, and V) and 4 nucleotides (A, T, G, and C), with a CatBoost model achieved an AUC of 0.980. This approach not only simplifies the feature generation process but also ensures efficient completion of high-throughput prediction tasks without compromising performance.

### Webserver Implementation and Standalone Application

We have developed a user-friendly web server and a python-based standalone application, which is publicly available to the scientific community. The web server is hosted at https://webs.iiitd.edu.in/raghava/lncrnapi, providing an interactive interface for users to submit lncRNA and protein sequences and predict LPIs without any computational setup. For more advanced users, we have developed Python-based standalone that can be downloaded from the webserver or our GitHub repository. We have also released lncrnaPI as a Python package on the Python Package Index (PyPI), facilitating the use of our tool directly in Python-based workflows and enabling seamless integration into computational pipelines for large-scale lncRNA–protein interaction prediction.

## Discussion

In this study, we developed a comprehensive set of computational models to predict lncRNA– protein interactions (LPIs), achieving results that clearly surpass existing tools. Traditional similarity-based methods performed poorly due to the limited sequence similarity between lncRNAs and proteins, leading to low accuracy and poor coverage. In contrast, our approach— combining the powerful CatBoost algorithm with contextual embeddings from large language models—proved remarkably effective. The final model achieved an MCC of 0.915 and an AUC of 0.989, setting a new benchmark in LPI prediction and demonstrating how advanced machine learning paired with modern sequence representations can capture the intricate nature of molecular interactions.

Our analysis of feature representations revealed a distinct hierarchy in predictive power. Basic compositional features like percentage composition of Leucine and Glycine were able to predict LPIs to a certain extent, achieving a maximum accuracy of 0.716. However, the best performance came from embeddings generated by DNABERT-2 and ESM2, emphasizing that the contextual knowledge encoded by large language models provides a much richer and more informative representation than traditional handcrafted features. While simple features were still effective, the LLM-based embeddings ultimately offered a more powerful way to represent biological sequences.

We also compared various modelling architectures, exploring both feature-based and end-to-end deep learning approaches. Although CatBoost delivered the top performance, our Siamese-like CNN achieved comparable accuracy (MCC - 0.908, AUC – 0.987) without any manual feature extraction, making it a compelling alternative. In contrast, alignment-based methods failed to predict LPIs accurately, highlighting the fact that LPI prediction depends on complex sequence patterns rather than simple homology, and that advanced machine learning techniques are essential to uncover these relationships.

Benchmarking our model against existing tools revealed major limitations – most of the methods rely on outdated or incomplete datasets (often NPInter v2.0) and are distributed only as raw source code, without accessible interfaces. In fact, we found only one functional tool, LPI-BLS, which performed no better than random on our updated dataset (AUC = 0.512). In contrast, our model, trained on the most comprehensive and up-to-date dataset, not only delivers superior predictive performance but also bridges the gap in usability. To make it broadly accessible, we have released lncrnaPI both as a user-friendly web server and a standalone Python package on PyPI, allowing seamless integration into computational pipelines and everyday research workflows.

## Conclusion

In this work, we developed and validated a new computational method that sets a higher standard for predicting lncRNA–protein interactions (LPIs). By combining modern machine learning algorithms with contextual embeddings learned from large language models, and by training on the most comprehensive and up-to-date dataset to date, our model delivers exceptional accuracy. It clearly outperforms the only existing functional tool and tackles long-standing issues in the field, such as the use of outdated datasets and the lack of accessible software. To make our approach widely usable, we’ve made the model available both as an easy-to-use web server and as a standalone Python package. We hope this resource will help researchers explore the complex regulatory roles of lncRNAs more effectively and accelerate discovery in this area.

## Funding source

The current work has been supported by the Department of Biotechnology (DBT) grant BT/PR40158/BTIS/137/24/2021.

## Conflict of interest

The authors declare no competing financial or non-financial interests.

## Acknowledgements

Authors are thankful to the Department of Science and Technology (DST-INSPIRE), Council of Scientific & Industrial Research (CSIR) and Indraprastha Institute of Information Technology, New Delhi, for fellowships and financial support. Authors are also thankful to Department of Computational Biology, IIITD New Delhi for infrastructure and facilities.

## Authors’ contributions

SC collected and processed the datasets. SC implemented the algorithms and developed the prediction models. SC, NB, and GPSR analyzed the results. SC created the front-end and back end of the web server. SC, NB and GPSR penned the manuscript. GPSR conceived and coordinated the project. All authors have read and approved the final manuscript.

## AUTHOR’S BIOGRAPHY

1. Shubham Choudhury is currently working as Ph.D. Student in Computational Biology from Department of Computational Biology, Indraprastha Institute of Information Technology, New Delhi, India.
2. Nisha Bajiya is currently working as Ph.D. Student in Computational Biology from Department of Computational Biology, Indraprastha Institute of Information Technology, New Delhi, India.
3. Gajendra P. S. Raghava is currently working as Professor and Head of Department of Computational Biology, Indraprastha Institute of Information Technology, New Delhi, India

## Notes

https://webs.iiitd.edu.in/raghava/lncrnapi/

### Competing Interest Statement

The authors have declared no competing interest.

